# Captive-bred Orangutans voluntarily choose to reward themselves with cabbage containing greater amounts of anthocyanin in a two-alternative decision task

**DOI:** 10.1101/099432

**Authors:** Neil Mennie, Rachael C. Symonds, Mazrul Mahadzir

## Abstract

Anthocyanins are an important part of the human diet and the most commonly consumed plant secondary metabolites. They are potent antioxidants, and in several recent studies the ingestion of anthocyanins has been linked to positive health benefits for humans. Here, we show that when given a choice between two alternative samples of cabbage to ingest, captive born orangutans (n = 6) voluntarily chose the sample that contained greater amounts of anthocyanin. This occurred when they had to decide between samples of red cabbage (*Brassica oleracea var. capitata f. rubra*) from the same plant (p<0.05), and samples from green cabbage (*Brassica oleracea var. capitata*) (p<0.01). This indicates that anthocyanin holds a reward value for these hominids. There was no difference in L*a*b* colour between ingested and discarded samples in red cabbage, but when the choice was between two green samples, the animals chose samples that were more green and yellow. There was also no difference in the amount of lightness (L*) between chosen and discarded samples of either plant. It is therefore unclear if the animals use leaf colour in decision-making. In addition to other macro nutrients provided by plants, anthocyanin is also chosen by these endangered apes.

## Introduction

Several fields of research in the behavioural sciences place an emphasis on the importance of reward. An early definition of a reward is an object that produces a change in behaviour (1), and that is *learning*. Rewards are considered objects, events, situations or activities that confer positive motivational properties via internal neural processes (2,3). Strongly associated with dopamine activity in midbrain structures such as the mesolimbic pathway and basal ganglia, rewards act as positive reinforcers by increasing the frequency of behaviours providing that reward. Rewards are important in aspects of learning, in approach behaviour, decision-making and economic models, addiction studies and in emotions (4). Rewards are also critical for actions that facilitate survival such as feeding and reproduction (2,5,6). As there is no specific sensory system dedicated to reward *per se*, the value of a reward needs to be calculated internally at the neuronal level (7,8). Such cost/benefit decision-making variables rely on inputs from the senses, and what aspects of the environment that are processed by the visual systems of animals to guide actions and goals will depend on their reward value (3,9,10).

Evidence suggests that colour vision evolved (sometimes convergently) in different animals to locate food rewards such as fruit from amongst a background of leaves or for identifying the plumage of conspecifics for reproduction (11–14). It’s also been suggested that trichromatic colour vision in primates has evolved to locate young edible leaves (15,16). Photoreceptors for colour would not have evolved without the presence of a concomitant signal, and using colour as a visual signal is mutually beneficial for plants and advantageous enough for them to have invested energy in producing such signals of reward. Fruit colours have been shown to be an adaptation to increase detectability of fruits for animals that disperse their seeds (17–19) and to assist the plant in inter-specific competition for pollinators and seed dispersers (20,21).

In addition to signalling macro-nutritional rewards such as carbohydrates and protein (19), there is also evidence that plants use signals that indicate *specific* nutritional reward to attract animals. The most important pigments that signal colour in ripe fruits are chlorophyll, carotenoids and anthocyanins (22). It is the combination of these different plant pigments that give flowers, fruit and leaves their distinctive colours (23).

The importance of carotenoid pigments as a pollinator attractant was demonstrated by researchers who introduced the YUP locus for carotenoid pigmentation reciprocally from a red *Mimulus cardinalis*, pollinated by humming birds into a pink *M. lewisii* pollinated by bees. The resulting transgenic *M. Lewisii* flowers were visited by humming birds and less frequently by bees, whilst the transgenic *M. cardinalis* flowers were now visited by bees as well as humming birds (20).

Anthocyanins are vivid blue, purple and red plant pigments widely distributed in fruits, flowers, leaves, stems and certain storage organs (24,25). They are members of the flavonoid group of compounds and interact exclusively with the visible spectrum between 500 and 550nm (26). Along with other flavonoids they have a role in protecting plants from various biotic and environmental stressors (27) and in animal attraction for the promotion of seed dispersal (17). Anthocyanins also act as antioxidants in plants both in vitro (28,29) and in vivo (30). The use of pigments such as anthocyanin by plants as a visual cue to attract animals and insects for pollination (20,21) or seed dispersal is well documented (18,19). Hoballah and colleagues (2007) looked at the role of anthocyanin as a visual attractant in pollination by introducing a gene encoding for an anthocyanin biosynthesis transcription factor from a pink flowered *Petunia integrifolia* to a white flowered *P. axillaris*. When offered a choice between white wild type flowers and pink transgenic flowers, hawkmoths chose white flowers and bumblebees chose pink flowers (31). Clearly the plant benefits from the use of this signal, but do the animals? Blackcaps show no colour preference when selecting fruit but showed a distinct preference for green over red insects, demonstrating that these birds show colour preference for their food depending on whether it is fruit or insect based (32).

Health benefits associated with a diet high in anthocyanins have been reported in both humans and animals (33). Several studies have highlighted an inverse correlation between a diet high in anthocyanin and a reduced prevalence of diseases such as cancer and heart disease (34). In addition, dietary anthocyanins have been reported to demonstrate antioxidant and anti-inflammatory properties. Rats with induced oxidative stress, to mimic metabolic syndrome, showed a marked improvement in antioxidant status when their diet was supplemented with Black Chokeberry extract containing a condensed form of anthocyanin glycosides (35). However despite this association between a diet high in antioxidants and a reduction in the incidence of certain diseases, this link has yet to be conclusively confirmed in animal or human clinical trials (36).

Anthocyanin is also linked to cognitive benefits. An analysis of the brains of aged rats fed on blueberry found that several anthocyanins were able to cross the blood-brain barrier into central areas such as the cerebellum, cortex, hippocampus and striatum. The aged rats also performed better than controls in a water maze task, suggesting learning and memory is facilitated by anthocyanin (37,38). What is not clear from that study is would the rats, actively seek out this reward if anthocyanins are to be considered a positive reinforcer that changes behaviour. Additionally, anthocyanins are odourless (39) and it remains unclear how such a reward could be sensed.

Other studies have documented the ability of birds, which possess tetrachromatic vision and have the ability to detect ultraviolet (UV), to distinguish food based on its colour (40–42). Using an avian eye model (12), Schaefer et al., 2008 studied the colour and pigment content of 60 different fruits consumed by birds, concluding that birds were able to discriminate fruits based on anthocyanin (but not carotenoid) content as fruits rich in anthocyanin are purple/black and reflect UV. They also noted that European blackcaps (*Sylvia atricapilla*) preferred food pellets soaked in anthocyanin to food that was not. Additionally, they found that migrating free-living birds preferentially selected fruits with a high anthocyanin/polyphenol content and this was postulated to protect against the oxidative stresses associated with migration (42). In addition to anthocyanins, studies have also documented attraction to the bright red, orange and yellow carotenoids. Cedar wax wings preferentially select red fruit over blue, yellow and green, (40) whilst the Great Tit, *Parus major* exhibited a preference for mealworms artificially enriched with carotenoids (43). However, despite anthocyanins comprising the largest group of plant pigments with over 600 isomers, there has been little exploration of the relationship between food anthocyanin content and visual selection and to our knowledge none in primates. Unlike birds, primates are trichromats and are unable to process wavelengths in the UV range. It therefore remains unclear as to whether they would actively seek food containing a greater anthocyanin content as a reward.

As part of a series of studies on the visual cognition of captive bred orangutans at Zoo Negara, the research documented in this manuscript addresses this question. Zoo, pet, laboratory and farm animals need continuous behavioural *enrichment* to maintain physical and cognitive capabilities (44–46), and for great apes this includes puzzles and problem solving tasks. As part of their enrichment, we gave the orangutans a simple decision making task. Our aim was to see if captive bred orangutans, when given a choice between two alternative samples of a plant that contain differing amounts of anthocyanin, voluntarily choose the sample containing greater concentrations of this flavonoid. Secondly, using a spectrophotometer, we measured the L*a*b* colour characteristics of these samples to see if there were any detectable visual differences between the samples rejected and those chosen for consumption that might be a visual signal of a reward to these endangered apes.

We simultaneously presented two identical-sized pieces of an everyday food source (cabbage) to the animals (n=6) using plastic kitchen spatulas and allowed the animals to choose ONE (the *winning* choice). The samples could either come from red cabbage (*Brassica oleracea var. capitata f. rubra*) or green cabbage (*Brassica oleracea var. capitata*). Green cabbage is part of the animals’ daily diet at the Zoo, while the red cabbage was chosen as it contains a greater amount of anthocyanin and is readily available throughout the year. Red cabbage is a fresh vegetable crop originally used as a medicinal treatment for headaches, gout and peptic ulcers (47). It has high levels of anthocyanin (48) with a complex content of up to 24 different anthocyanins currently identified (49,50). This high level of anthocyanin has been positively correlated with total antioxidant content (51). Red cabbage anthocyanin extract has also been shown to reduce the incidence of colorectal cancer in male rats fed a diet of red cabbage colour natural extract (52).

As green cabbage is part of the animals’ daily diet, then should we find a preference between two green samples for the one containing higher concentrations of anthocyanin, we cannot be sure that this was not due to long-term learning. We therefore also need to test a relatively novel, safe food source to remove this confound. All orangutans had previously encountered red cabbage, but it is never part of their daily diet as the costs are prohibitive. This makes it ideal to test alongside the green, for it contains greater amounts of anthocyanin than the green and is not encountered by the animals daily. However, should the animals prefer red cabbage instead of green, then this could be due to either stimulus novelty (53) or because they pay greater visual attention to red food (54,55) and not necessarily due to the higher levels of anthocyanin within those red/purple leaves. Accordingly, we split the samples into high and low anthocyanin groups and then tested six different combinations of samples: Red High/Red Low, Red High/Green High, Red High/Green Low, Red Low/Green High, Red Low/Green Low and Green High/Green Low. We looked for a preference of the animals’ for samples containing higher amounts of anthocyanin and specifically within “same-colour” combinations to rule out the role of novelty and to see if colour was a factor. A further advantage of varying the colours is that this provides greater enrichment during the task for these highly intelligent animals.

## Methods

### Participants

Three female Bornean orangutans (*Pongo pygmaeus*), and three Sumatran orangutans (*Pongo abelii*) (1 male, 2 female) participated in this study. Five animals were adult, with one juvenile (a female Sumatran, aged 12). Ages ranged from 12 to 26 years old (Ave. 19). All animals undergo regular health checks by the veterinary staff at Zoo Negara and were in good health throughout the study.

### Ethics

This study was conducted with the assistance of veterinary staff and Zookeepers at Zoo Negara as part of the Zoo’s *enrichment* programme. It was also conducted with the full permission of the Wildlife Department of Peninsular Malaysia (Perhilitan) as part of a series of behavioural studies run by Dr Neil Mennie of the University of Nottingham Malaysia Campus and with the approval of the Faculty of Science at the University of Nottingham Malaysia Campus.

### Apparatus and Materials

Cabbage leaf preparation and leaf pigment determination

The varieties of cabbage used were Red cabbage (*Brassica oleracea var. capitata f. rubra*) and green cabbage (*Brassica oleracea var. capitata*). Cabbage leaf material was purchased from a local supermarket on the day prior to each zoo visit and prepared on that day. Cabbages were ensured to be of the identical type and from the same supplier. Cabbage leaves were cut into polygonal 5-pointed star shapes using a standard cookie cutter (area ~0.3cm^2^). The thickness of each sample varied according to the individual leaf thickness. See fig.1 for images of cabbage samples. Samples were kept refrigerated in airtight plastic bags prior to transportation to the zoo.

Leaf SPAD chlorophyll and anthocyanin content were measured non-destructively using a portable Minolta Chlorophyll Meter SPAD-502 (Konica Minolta, Langenhagen, Germany) and an ACM-200 Plus Anthocyanin Content Meter (Opti-Science, Inc. Hudson, New York, USA), respectively. Three readings were taken per sample and averaged to give a final value.

Cabbage leaf colour was assessed using a quantitative three dimensional (X, Y & Z axis) colour method (Commision Internationale de l’Eclairage 1976 or CIE (L*a*b)) using a Konica Minolta CM-600d Spectrophotometer. To measure individual leaf colour, three replicate measurements were taken and averaged to determine CIE L*a*b.

The star shaped cabbage pieces were sorted into four significantly different groups (p<0.001) based on their anthocyanin content value: red leaf high anthocyanin (RH), red leaf low anthocyanin (RL), green leaf high anthocyanin (GH) and green leaf low anthocyanin (GL). These 4 groups were paired with each other to produce six pairings: (1) red high & red low (RH/RL), (2) red high & green high (RH/GH), (3) red high & green low (RH/GL), (4) red low & green high (RL/GH), (5) red low & green low (RL/GL) and (6) green high & green low (GH/GL). See fig. 1. A total of 788 red cabbage leaves (394 RH & 394 RL) and 780 green cabbage leaves (390 GH & 390 GL) were measured for stimuli presentation in this study.

The samples were presented to the animals at their cage door using two identical plastic kitchen spatulas, consisting of a grey handle and black spoon (see fig. 1 and video.1 below) on the following day. The time between recording leaf pigment values and presentation to the animals was never more than 24hrs. All spatulas were washed and sterilised between sessions, with different spatulas used on each animal to minimise disease transmission.

**Figure 1:**
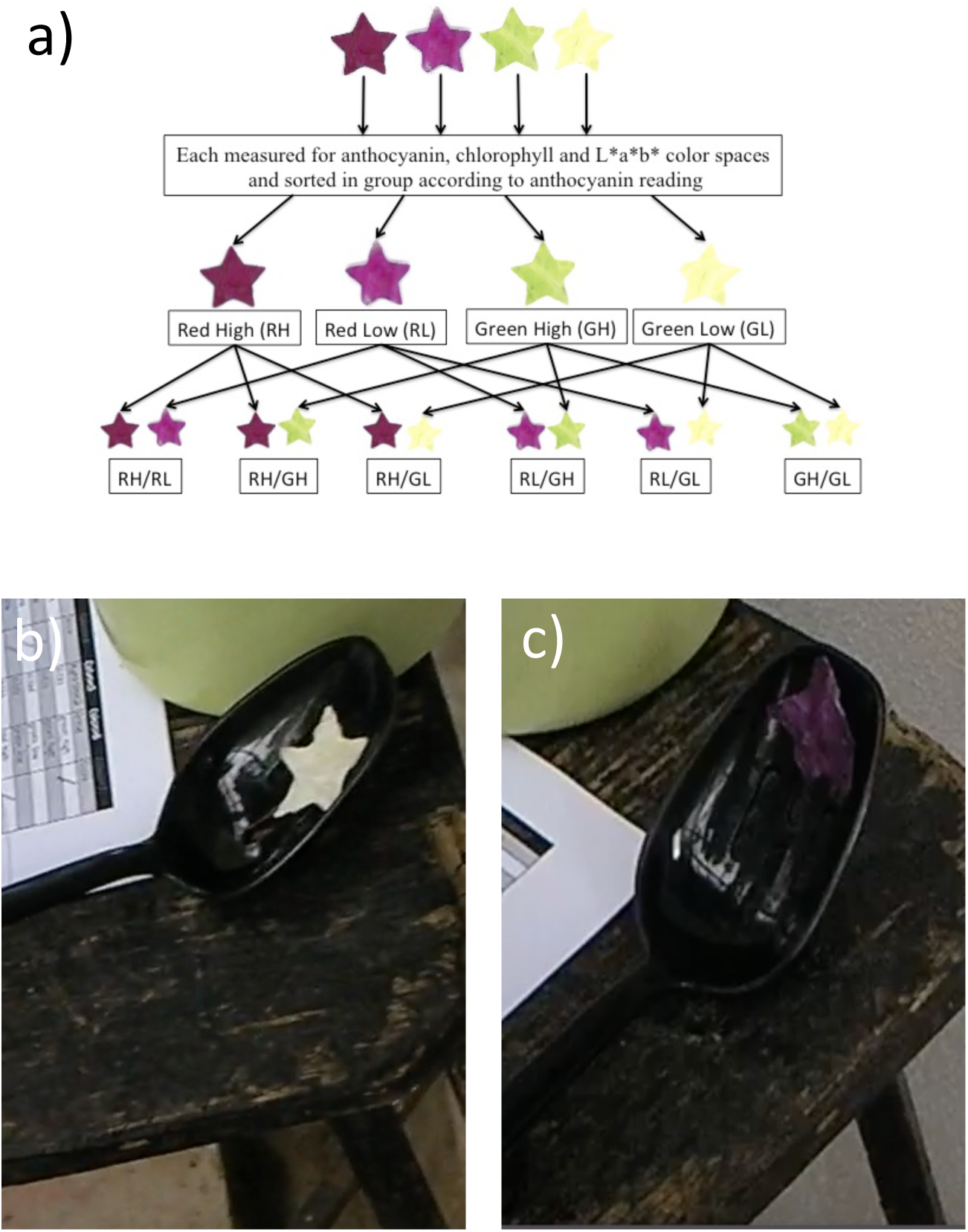
*Cabbage samples*. Figl.a Red and green cabbages are sorted into groups according to high and low anthocyanin readings. These 4 groups are then combined to give 6 different pairings that were presented to the Orangutans on the subsequent day. Fig1.b and fig1.c are images of green and red samples of cabbage on a spatula.

### Food-Choice Procedure

1. Rather than using pre-determined thresholds for high and low anthocyanin content, we split the samples into high and low groups each time the samples were prepared in the lab. After cutting and determining leaf pigment values of 60 samples (30 red & 30 green), the samples were then sorted into the 4 different groups (RH, RL, GH, GL). As pigment values vary widely from plant to plant, this means that a particular sample with a specific anthocyanin reading that fell into a “high” group on one day could subsequently be classed as a “low” reading (and vice versa) on another day. However, in practice this rarely occurred. Samples were prepared twice a week.
2. The samples were stored overnight in a standard refrigerator (4°C), and taken to the Ape-centre at the National Zoo on the following morning, where they were sorted into the 30 pairings and then placed in a plastic folder in trial order.
3. On each visit to the ape-centre, two different animals were each presented with 15 randomised pairings (15 trials) using a black kitchen spatula in each hand of the experimenter.
4. A small table was placed in front of the cage door. On that table were the 2 identical spatulas, pen and paper to record the animal’s choice, and a green plastic “trash can” where the unselected samples from each trial would be deposited. The folder containing the cabbage pairs was situated on a second small table.
5. For each trial, the experimenter removed a pair of samples from the folder, placed them on the spatulas in front of him, and then simultaneously raised the spatulas to the cage door where the animal made its choice through the bars. The experimenter also ensured red or green samples appeared equally frequently in either hand, and randomly employed switch trials – where during presentation of the stimuli s(he) crossed their arms. This serves two purposes; it ensures the animal maintains attention on the task and it also allows us to easily detect any Left or Right bias the animal might show for either hand of the experimenter *or in their own reaching and grasping movement*. The accompanying video shows 3 successive trials, including a switch trial, to a male Sumatran Orangutan: (see supplementary video).
6. After every 5 trials, the animal was presented with a reward in either L or R spatula. The reward was a grape (red or green), or a red or yellow cherry tomato. This limits the effects of boredom that highly intelligent animals might suffer when doing this task and encourages them to attend to the experimenter. Average time to complete the 15 trials (and 3 reward trials) was approx. 15 min.

### Training Procedure

Prior to experimental trials, orangutans were familiarised with the experimental paradigm with twice-weekly training visits over 3-6 weeks depending on the animal. During training, they were given the choice of a number of differing fruit, vegetables and nuts from the spatulas after first learning to take one item of food from one spatula. Once this was learnt (within 2 sessions), they then had to learn to choose between one or two random food items presented in parallel. All animals initially tried to grasp *both* items of food, either simultaneously using both hands, or serially using one after another. To train them that they could only receive their first choice, the spatula on the unselected side was withdrawn as a soon as any discernible reach to the preferred side was in evidence. That sample was then placed in a green “trash can” in view of the animals. Video1 shows the experimenter withdrawing the spatula from the less-preferred side and depositing that sample in the “trash can”. Animals were deemed to have learnt the task once they had ceased to make any reaches for the second spatula and continued to do so for 3 successive weeks. If an animal showed little or no interest in participation in training sessions or in later experimental trials, they were not disturbed and the experimenter went on to work with another animal.

### Control Readings of Leaf Samples

The aim of this study was to determine whether Orangutans choose to consume one of two alternative pieces of cabbage based on the levels of anthocyanin or chlorophyll present in those samples. However, as levels of anthocyanin in a leaf are correlated with leaf colour, it is hard to tell if they are making a choice based on the colour of that leaf, irrespective of anthocyanin content. It’s important that we record any potential changes in leaf pigment values and spectrophotometer readings that might occur over the 24hr period between preparation and consumption to see how they differ when presented to the animals. Therefore, after all initial readings were taken we stored 10 red and 10 green samples in the folder in a refrigerator at 4°C. We took further readings 1 and 3 days later. These values are shown in the results.

### Statistical analysis

All data is reported as mean values with a standard error (SE). Winning and losing cabbage choices were analysed using within subjects T-test. The anthocyanin content of the four cabbage groups was analysed by one-way Analysis of Variance (ANOVA) and pairwise multiple comparisons were determined using Tukey’s Pairwise Comparison test (P>0.05). Correlation coefficients were used to analyse anthocyanin, chlorophyll and L*a*b relationships using Genstat 16th edition.

## Results

### 1.1 Do levels of anthocyanin and chlorophyll change over the 24hrs between preparation and consumption?

**Figure 2:**
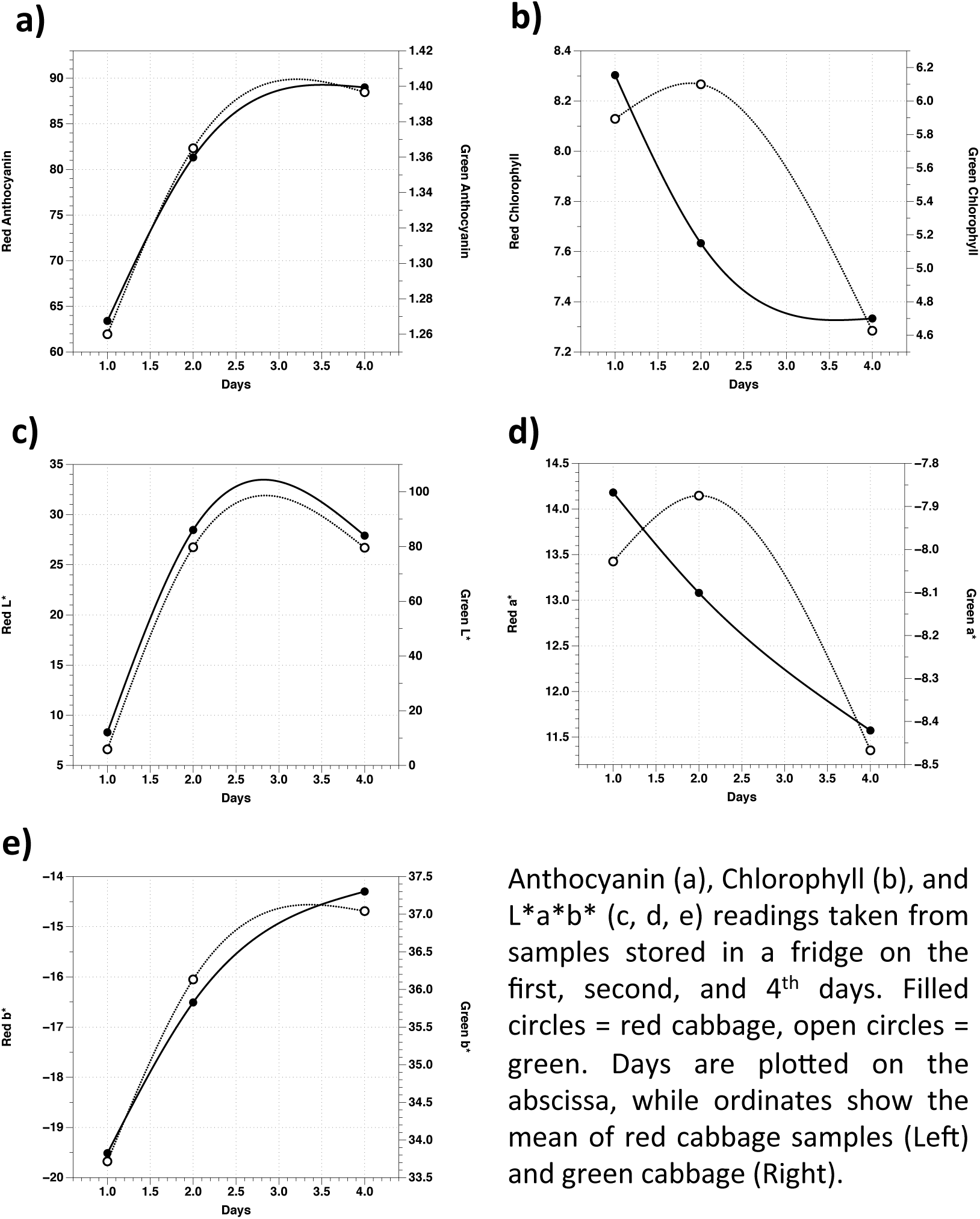
*Changes in Anthocyanin, Chlorophyll and Spectral measurements of red and green cabbage over 3 days*.

Figure 2 illustrates mean values of red (n=10) and green (n=10) samples of cabbage that were stored for 4 days in the fridge, with readings taken on day 1 (the day of preparation), day 2 (24hrs later) and on day 4. We did not separate our control samples into high and low conditions. All plots have been smoothed. Levels of anthocyanin increased in *both* types of cabbage over 24hrs in a dark fridge (fig. 2a), indicating that the animals were consuming cabbage samples that contained higher levels of anthocyanin than that recorded on the previous day. Red cabbage shows an increase in anthocyanin (63 to 81), which is a 28% average increase. Green cabbage shows a smaller increase (1.26 to 1.37), which is an average 8% increase in anthocyanin levels over the 24hr period of interest. Two days later (day 4), this increase is smaller (9% red and 2% green). Note that the overall readings are very different for the two types of cabbage, with red cabbage plotted on the left ordinate and green on the right. Anthocyanin levels do not decrease as much in the dark as in the light (56), and our findings suggest that as both red and green cabbage show an increase in anthocyanin then any differences between preparation and consumption by the animals should not influence our results if the animals are making a choice based on anthocyanin levels.

As for chlorophyll, amounts of this molecule *decreased* in red cabbage by 8% over 24hrs, while the green shows a small increase of 4% (see fig. 2b). Figure 2b shows that on day 2, red cabbage decreased from 8.3 to an average reading of 7.6, while green cabbage values rose from 5.9 to 6.1. Chlorophyll values for the two types of plant are closest on day 2, which is equivalent to the time of consumption by the animals, and seems to be mainly due to a decrease in the chlorophyll content of the red cabbage with the green remaining fairly constant. It is unlikely that when the animals consume the samples after 24 hours that chlorophyll levels will be different.

### 1.2 Do spectral readings change over the 24hrs between preparation and consumption?

The remaining plots in figure 2 (c,d,e) illustrate the concomitant spectral readings taken over the 4 days. If the animals are making a decision based on visual cues, then cues such as lightness (L*) or red/green (a*) of the food are an important factor in food choice (20). Our data shows that L* in fig. 2c closely corresponds to the anthocyanin readings in fig. 2a, with an increase over 24hrs for both types of cabbage before levelling off. Green is clearly much lighter than red, but the rate of increase in lightness is not different. As this affects both types of cabbage, then it should not adversely affect our results.

We also see a decrease in redness of red cabbage, with readings moving further towards the green in the red/green channel (a*), see fig. 2d. Additionally, as the chlorophyll decreases in red cabbage, we see an increase in yellowness in the blue/yellow channel (b*) for the red. For green cabbage it is different. Spectral readings of green cabbage samples show that yellowness increases with a decrease in chlorophyll (as they do for red cabbage, fig. 2e), but they differ from the red cabbage in that green cabbage becomes less green and redder (fig. 2d).

### 2. How is anthocyanin distributed in red and green cabbage?

Before looking to see if the animals prefer samples of cabbage with differing anthocyanin levels, we first need to see if our arbitrary separation of all samples into 4 groups each week actually resulted in significantly different levels of anthocyanin. Table 1 illustrates the overall anthocyanin values recorded from the four differing groups of red and green cabbage, while figure 3 shows the distribution of these samples.

**Table 1:** *Overall Distribution of Anthocyanin means in different samples of cabbage*.

|  | **Green Low** | **Green High** | **Red Low** | **Red High** |
| --- | --- | --- | --- | --- |
| **Mean** | 1.02 | 1.21 | 39.21 | 88.50 |
| **Median** | 1.00 | 1.20 | 33.97 | 84.57 |
| **Mode** | 1.00 | 1.10 | 29.67 | 62.80 |
| **Standard Deviation** | 0.08 | 0.13 | 21.03 | 36.49 |
| **Standard Error** | 0.00 | 0.01 | 1.06 | 1.84 |
| **N** | 370 | 368 | 394 | 394 |

**Figure 3:**
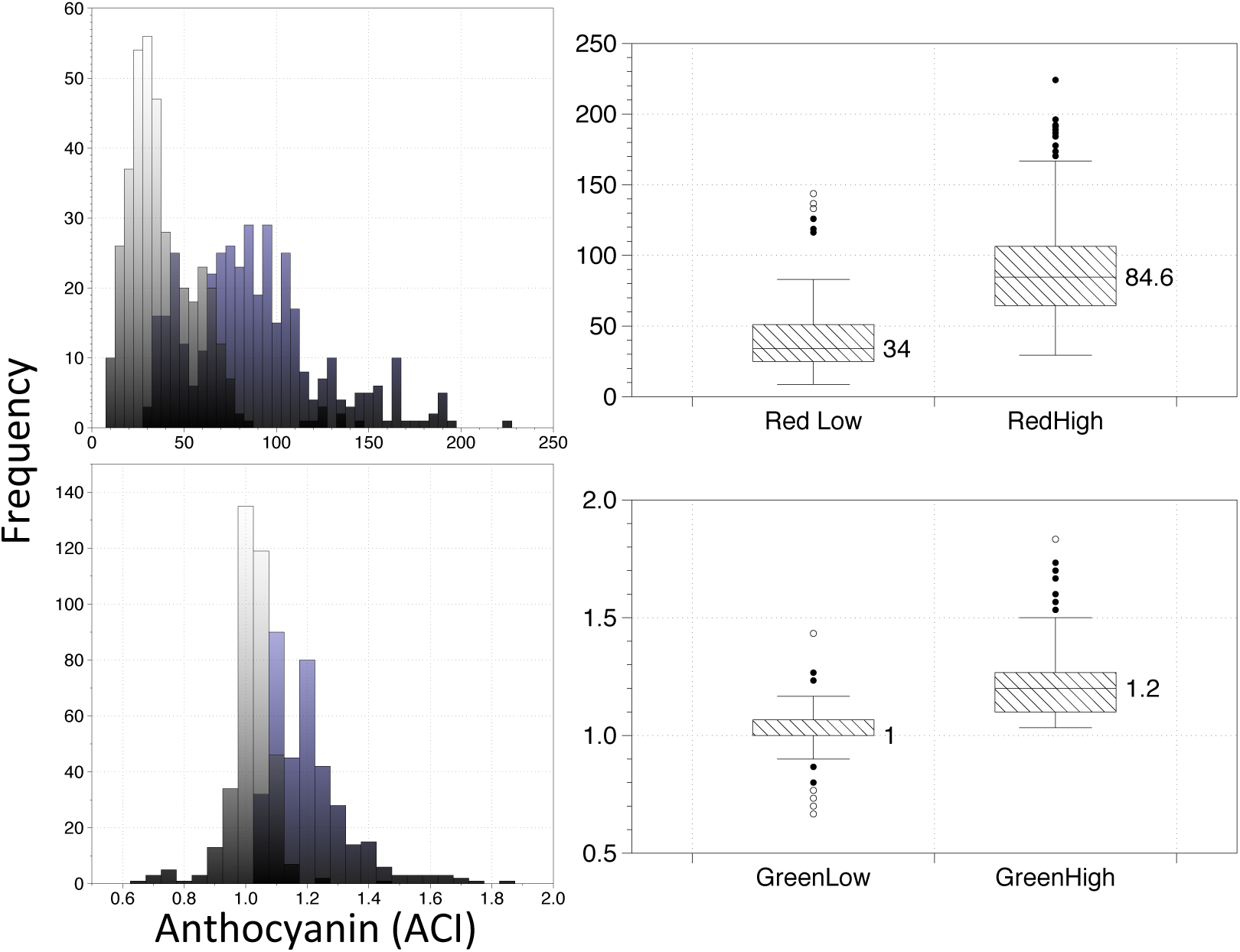
*Distribution of Anthocyanin in different samples of cabbage*. The top row shows a histogram of the frequency of all red high (purple) and red low (grey) samples on the left while the box plots on the right illustrate the median, interquartile ranges and outliers of these two groups. The frequency of samples of Green High (purple) and Green Low (Grey) are illustrated in the bottom row, along with respective boxplots on the right.

As Table 1 illustrates, the overall mean anthocyanin values of red cabbage (88.5 versus 39.2) are clearly much higher than the green (1.2 versus 1). Additionally, the HIGH and LOW groups are significantly different to each (p<0.001). The level of anthocyanin in each of these 4 groups was different to each other prior to the animals’ choice and irrespective of which of the 6 pairings those samples were then subsequently assigned.

Histograms in fig. 3 show that there was some overlap so that, for example, if a sample was scored as LOW on any one day then the same sample might fall into the HIGH class on another day. Nevertheless, there was a sufficient difference between HIGH and LOW in both types of cabbage to show that in addition to clear differences in anthocyanin levels between red and green cabbage the HIGH and LOW groups within those separate plants are also different to each other.

To establish if levels of chlorophyll correlated with the levels of anthocyanin measured, correlation coefficients were carried out between these two pigments and spectrophotometer L*a*b* readings (Table 2). The results show that in both red and green cabbage leaves, anthocyanin was significantly positively correlated with chlorophyll (p<0.001). Increasing levels of anthocyanin were negatively correlated with L* (p<0.001), which correspond to lightness and b* (p<0.001) which corresponds to decreasing green-yellow and increasing blue. Levels of a* which correspond to increasing redness were positively correlated with anthocyanin (p<0.001).

**Table 2:** *Correlation coefficients (^2^) for anthocyanin, chlorophyll andL*a*b* in red and green cabbage leaves*.

|  |  |  |  |  |  |
| --- | --- | --- | --- | --- | --- |
| **L\*** | - |  |  |  |  |
| **a\*** | 0.66** | - |  |  |  |
| **b\*** | 0.90** | -0.68** | - |  |  |
| **Chlorophyll** | -0.62** | 0.27** | -0.37** | - |  |
| **Anthocyanin** | 0.78** | 0.43** | -0.63** | 0.77** | - |
|  | **L\*** | **a\*** | **b\*** | **Chlorophyll** | **Anthocyanin** |

### 3.1 Can Orangutans consistently choose the sample containing more anthocyanin?

The first choice of each animal was recorded and is shown as the WINNING column in table 3 below. The mean anthocyanin value of winning and losing choices are illustrated next to the 6 conditions the animals were presented with. The LOSING choice was discarded after each trial into a green bucket in view of the animal (See accompanying video). Individual mean values were calculated for each animal, and the mean of those values are presented in table 3. Table 3 also shows the significance levels of 6 separate within subject t-tests.

**Table 3:**
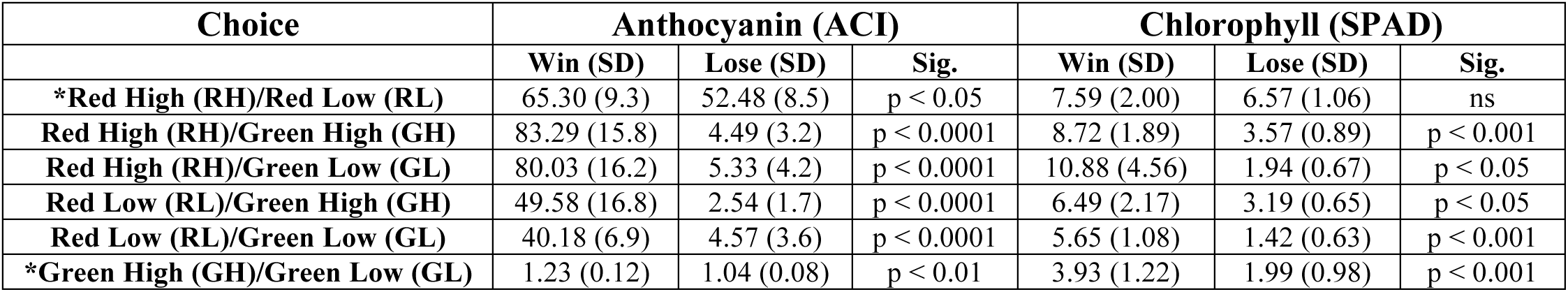
*Mean Anthocyanin and Chlorophyll levels in Winning and Losing Choices*. The table shows the mean values of anthocyanin and chlorophyll of the samples in each of the six conditions. Brackets = 1SD.

| Choice | Anthocyanin (ACI) |  |  | Chlorophyll (SPAD) |  |  |
| --- | --- | --- | --- | --- | --- | --- |
|  | Win (SD) | Lose (SD) | Sig. | Win (SD) | Lose (SD) | Sig. |
| **\*Red High (RH)/Red Low (RL)** | 65.30 (9.3) | 52.48 (8.5) | p < 0.05 | 7.59 (2.00) | 6.57 (1.06) | ns |
| **Red High (RH)/Green High (GH)** | 83.29 (15.8) | 4.49 (3.2) | p < 0.0001 | 8.72 (1.89) | 3.57 (0.89) | p < 0.001 |
| **Red High (RH)/Green Low (GL)** | 80.03 (16.2) | 5.33 (4.2) | p < 0.0001 | 10.88 (4.56) | 1.94 (0.67) | p < 0.05 |
| **Red Low (RL)/Green High (GH)** | 49.58 (16.8) | 2.54 (1.7) | p < 0.0001 | 6.49 (2.17) | 3.19 (0.65) | p < 0.05 |
| **Red Low (RL)/Green Low (GL)** | 40.18 (6.9) | 4.57 (3.6) | p < 0.0001 | 5.65 (1.08) | 1.42 (0.63) | p < 0.001 |
| **\*Green High (GH)/Green Low (GL)** | 1.23 (0.12) | 1.04 (0.08) | p < 0.01 | 3.93 (1.22) | 1.99 (0.98) | p < 0.001 |

All six animals preferred red cabbage to green, and the large differences in the mean values of anthocyanin between winning and losing choices in the 4 middle rows of table 2 reflect this. However, each occasionally chose a green instead of a red sample, and because of the large differences in the anthocyanin content of red and green (see table 1), this raised the mean of the overall losing choice to a level above that of green cabbage alone. This is illustrated in the losing value of the 4 middle rows in table 3. The 2 conditions where the animals were presented with samples of the same colour (* in table 3) are of more interest, as it is much harder to distinguish if any one sample contains a greater amount of anthocyanin without green and red cues.

### 3.2 Can Orangutans choose the sample that contains more anthocyanin when presented with two reds?

Yes. The first row in table 3 shows that, when presented with a choice of two red cabbage samples from the same plant, the winning choice contained a greater mean amount of anthocyanin (p<0.05). Figure 4 below illustrates this difference, along with individual boxplots (fig. 4b) and individual means (fig. 4c) for each animal. Five of the six animals consistently chose the cabbage with higher anthocyanin content (RED HIGH). Only one animal (Tsunami) more often chose the red sample with lower anthocyanin content. Overall, captive-bred Orangutans prefer red cabbage that contains more anthocyanin.

**Figure 4:**
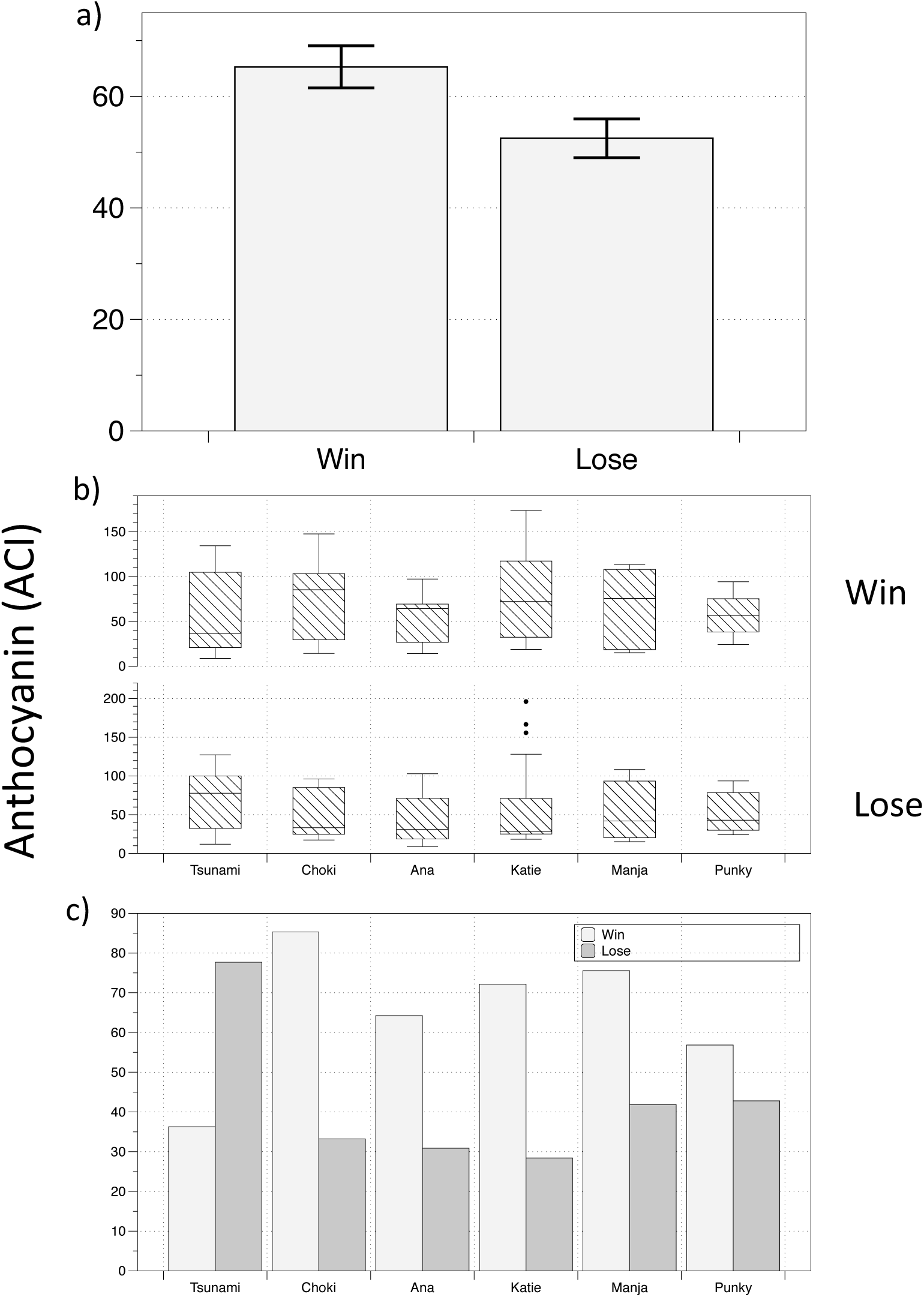
*Winning choices for Red High versus Red Low samples*. In a) the mean values of anthocyanin content for winning and losing choices is illustrated. Bars indicate +/- 1SE. In b) individual boxplots for each animal, illustrating interquartile ranges and median values. Dots indicate outliers. In c) the individual means are shown.

### 3.3 Can Orangutans choose the sample of green cabbage that contains more anthocyanin when presented with two greens?

GREEN HIGH/GREEN LOW choices are just as important. These are the samples with little anthocyanin present in either spatula. However, the RED HIGH/RED LOW choices are where large amounts of this odourless molecule are found and directly contribute to visual properties such as “redness” and “purple”. Perhaps, based on these visual properties, the animals might find it easier to discriminate between the red samples than the green, and so examining their ability to discriminate between green samples (low anthocyanin) is very necessary. Secondly, as the animals get a daily supply of green cabbage in their diet and are unable (or unwilling) to discriminate between green leaves, then it might suggest that RED HIGH/RED LOW results could be due to stimulus novelty, and not necessarily due to levels of anthocyanin.

Our captive-born Orangutans also consistently choose the green cabbage that contains more anthocyanin. Figure 5 shows that all 6 animals choose the sample of cabbage with higher anthocyanin content while the losing choice contained less anthocyanin. The mean values of winning and losing choices are illustrated in fig. 5a, while individual median values are shown in the boxplots of fig. 5b and individual mean values in fig. 5c. Their ability to discriminate between red samples isn’t just down to stimulus novelty, and the mean differences between the two green samples are of greater significance.

**Figure 5:**
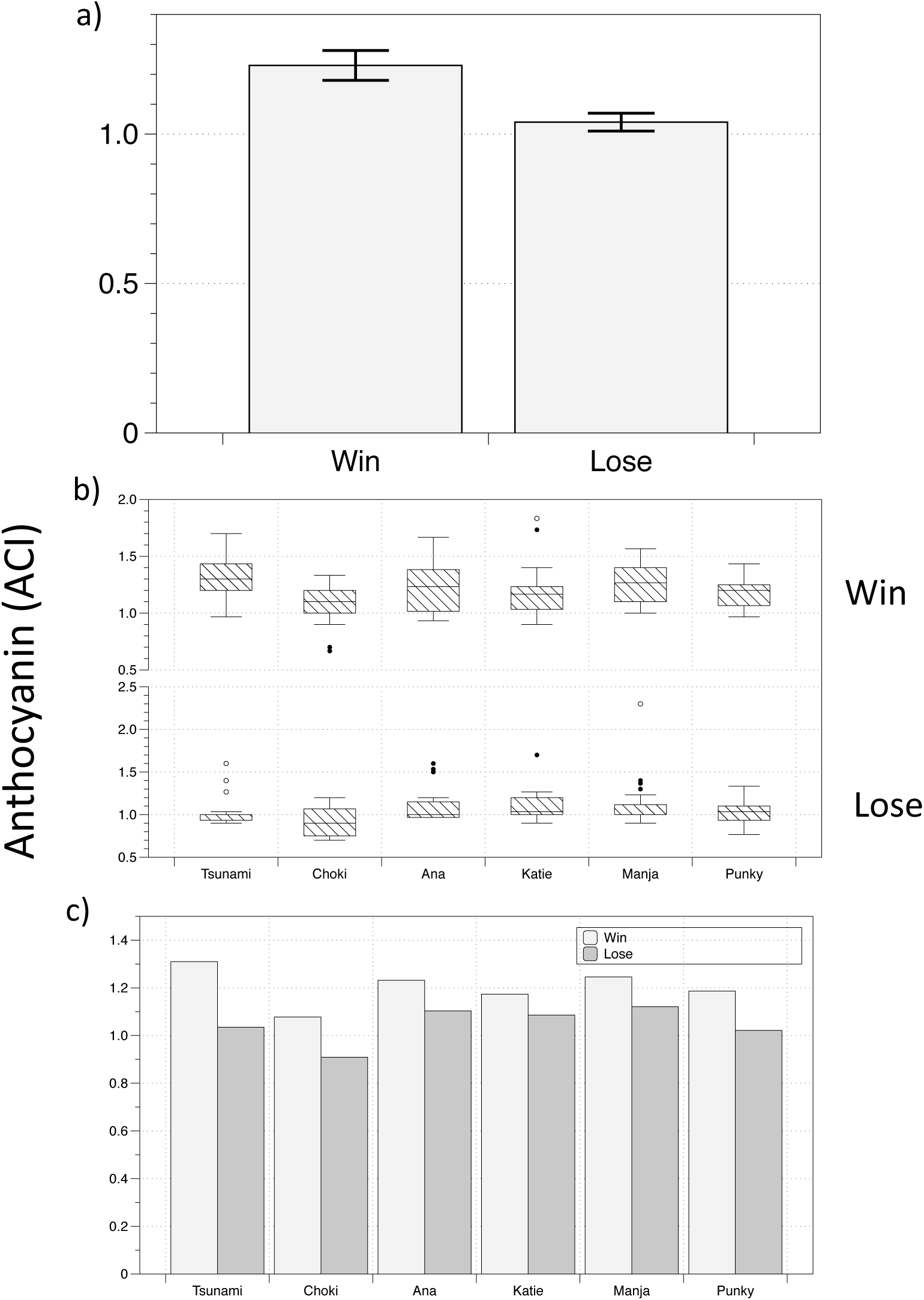
*Winning choices for Green High versus Green Low samples*. In a) the mean values of anthocyanin content for winning and losing choices is illustrated. Bars indicate +/- 1SE. b) Individual boxplots for each animal, illustrating interquartile ranges and median values. Dots indicate the outliers. c) Individual means.

### 4. Are the winning and losing choices different in colour or lightness?

The results clearly show that winning choices have a greater concentration of anthocyanin than losing ones. Additionally, our data also indicates that in the 4 conditions when an animal was given a choice between red and green cabbage, they almost always preferred red cabbage. However, as Table 4 (below) shows, the mean values of Lightness (L*) for these winning red choices were also significantly *darker* (see also figure 1). Table 4 also shows that for the two conditions where the animals had to choose between samples from the same green or red plant there was no significant difference in lightness. This is also illustrated in figure 6.

**Table 4:**
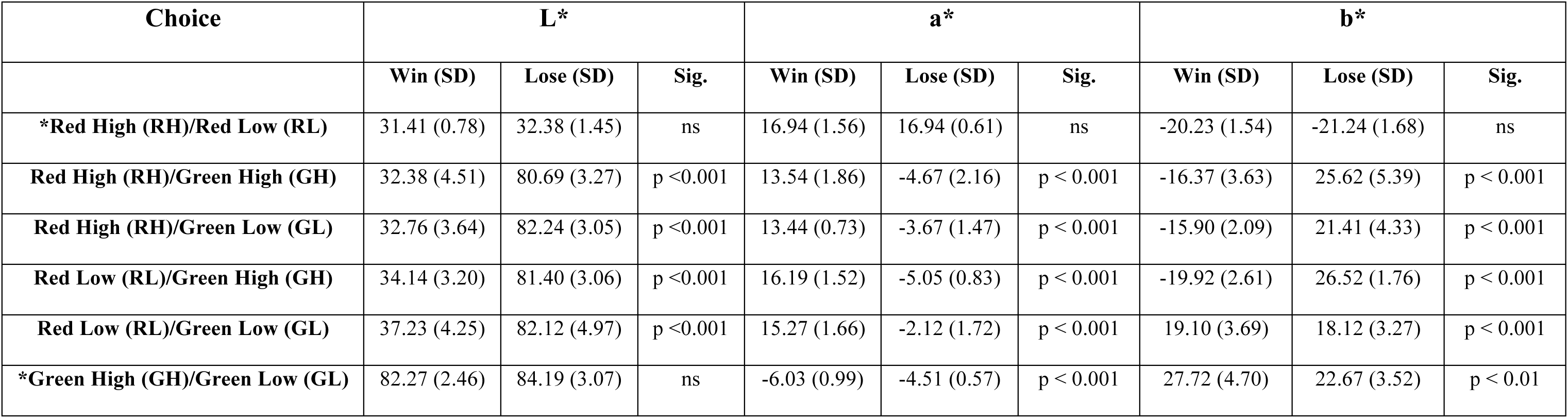
*Mean values of Lightness (L*), red/green (a*) and yellow/blue (b*) of the winning and losing choices in each of six conditions*. Brackets = 1SD.

The spectrophotometer provided 3 readings for each sample. First, the lightness, L* represents the darkest black at L* = 0, and the brightest white at L* = 100. The color channels, a* and b*, represent true neutral gray values at a* = 0 and b* = 0.The red/green opponent colors are represented along the a* axis, with green at negative a* values and red at positive a* values. The yellow/blue opponent colors are represented along the b* axis, with blue at negative b* values and yellow at positive b* values. Looking specifically at conditions where the animals had to choose between samples from the same plant, the data presented in figure 6 are the lightness (L*) readings for the winning and losing choices in the RED HIGH/RED LOW and GREEN HIGH/GREEN LOW conditions (see fig. 6a). In fig. 6b the mean values of winning and losing choices for these 2 conditions in the red/green opponent channel (a*) are shown, while fig. 6c illustrates the results for the blue/yellow (b*) channel of these 2 conditions.

**Figure 6:**
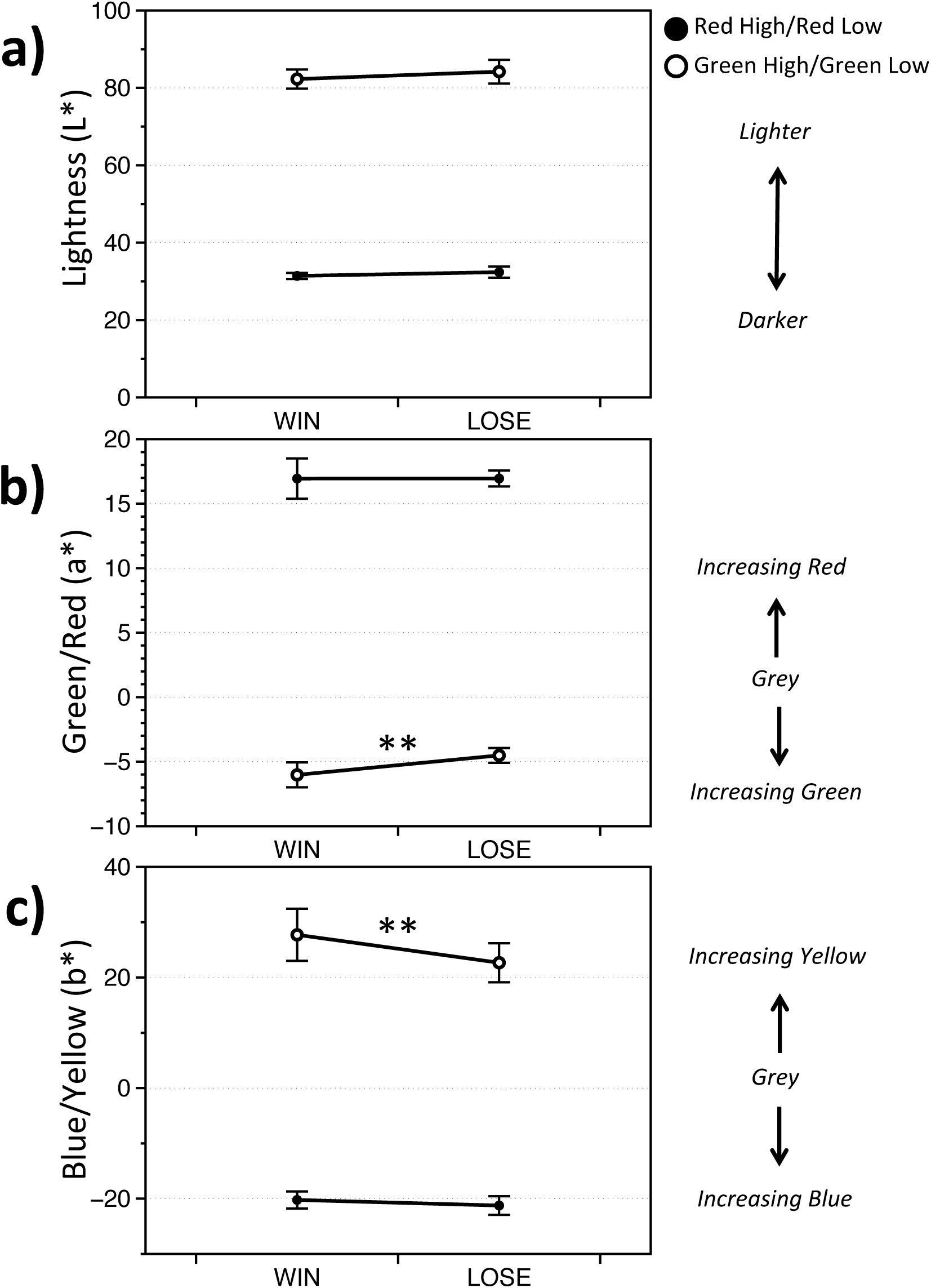
*Mean L*a*b* values of winning and losing choices of cabbage in the red high v red low and the green high v green low conditions*. Significant differences between the mean values are indicated by **. Neutral Grey is zero in b) and c).

Figure 6 and table 4 above show that there was no difference in the lightness (L*) of the samples that were consumed, i.e. the winning choice, compared to the losing choice in those two conditions. Fig. 6a shows that while green cabbages are clearly much lighter than red ones (also see photographs in fig. 1), there was no difference in lightness between WIN or LOSE in either RH/RL or GH/GL conditions.

Additionally, fig. 6 also shows that when choosing between two red samples, there was no difference between mean values of winning and losing choices along either the red/green channel (a*) or the blue/yellow (b*) channel. Fig. 6b and fig. 6c illustrate this, showing that even though the winning choice between two reds contained more anthocyanin, it doesn’t appear to differ in colour. Winning and losing choices in the RH/RL combinations are equally red and blue. When presented with the 4 “different colour” combinations (RH/GH, RH/GL, RL/GH and RL/GL) the winning choice was significantly more red than green (a*). These combinations were also more blue than yellow (b*) with the exception of RL/GL in which the willing choice was more yellow than blue (Table 4).

However, there *are* differences between winning and losing choices in the GH/GL condition (Table 4). Not only does the winning sample contain more anthocyanin, but it also appears to be significantly greener (Mean: −6.03, SD: 0.97) than the losing choice (Mean: −4.51, SD: 0.57) on the red/green opponent channel (see fig. 6b). It is also significantly more yellow. The mean value of the winning choice on the blue/yellow opponent channel in fig. 6c is 27.72 (SD: 4.70) while the mean value of the losing green choice was 22.67 (SD: 3.52). Therefore, the green samples that all animals chose to consume were just as dark as the ones that they rejected but winning choices were significantly more green and yellow. When it comes to an intra-green choice these differences in colour might be perceptible visual signals that were available to orangutan decision-making mechanisms. These differences are highlighted in fig. 7 below, where we have plotted mean a*, b* values for each animal in the “same-colour” conditions to illustrate their choices in colour space.

**Figure 7:**
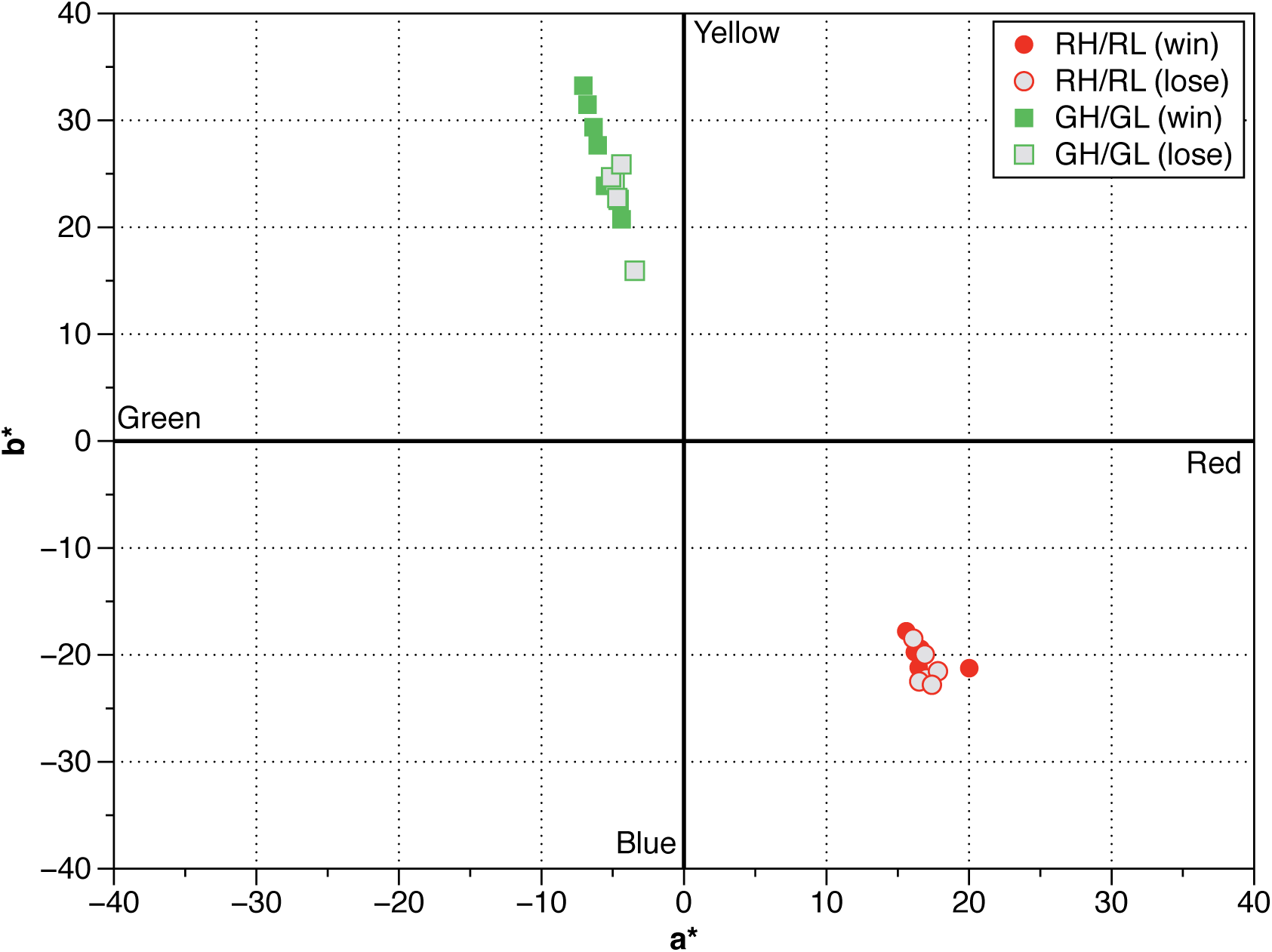
*Position of the a* and b* values for each Orangutan’s winning and losing choice in the RH/RL and GH/GL conditions*. a* is plotted along the x-axis and b* along the y-axis. Neutral grey is zero in the colour space.

### 5. Are orangutans making decisions based on the amount of chlorophyll in cabbage and not on the levels of anthocyanin?

As anthocyanin and chlorophyll have been previously shown to be correlated (57), it’s possible that the animals are making decisions on what to consume based on the chlorophyll content and not anthocyanin. Chlorophyll was also positively correlated with anthocyanin in this study and it is therefore hard to distinguish between these two alternatives (p<0.001). Table 3 shows that there were significant differences in the amount of chlorophyll between the winning and losing choices for all conditions except for the RH v RL condition (p = ns).This suggests that for red cabbage leaves that come from the same plant the animals are choosing based on anthocyanin rather than chlorophyll leaf content as there was a significant different between RH/RL (p<0.05) (Table 3). Nevertheless, we can not conclusively exclude the role of chlorophyll based on this data.

## Discussion

We set out to test two specific aims. Would captive orangutans choose the samples of cabbage that contained higher levels of anthocyanin? Secondly, as anthocyanin is odourless (39), could we identify any visual differences between winning and losing samples based on lightness (L*) or colour. This research is important for wild orangutan conservation strategies, management of captive orangutan populations, and in humans for vision science, food science and nutrition.

Our findings support the first objective. Six captive-bred Orangutans of two different species, *Pongo pygmaeus* and *Pongo abelii* (58–60) voluntarily chose two different types of cabbage containing higher levels of anthocyanin. This was not just a preference for red cabbage (which contains much higher levels of anthocyanin) that could be due to either the novelty of the stimulus or to an inability to ignore salient colours such as red/purple that attract attention (54,55). When the choice was between samples that both came from the *same* red or green plant, the animals consistently chose the sample containing more anthocyanin. Their decision is unlikely to be due to either saliency or novelty, and points to a preference for leaves that contain anthocyanin irrespective of the colour red.

Using a spectrophotometer to investigate the second hypothesis we found that there was no difference in the lightness (L*) between winning and losing choices. Lightness is an important component of colour and crucial for object identification, to discriminate between two objects and L* influences the *contrast* between the sample and the black background of the spatula This is again important in attracting visual attention (55,61,62). Not finding any differences in L* between winning and losing choices was therefore surprising. Perhaps this was because we could not control for levels of ambient background lighting *at* the moment of presentation and which could influence the L* perceived by the animal at decision time. Nevertheless, orangutan decision-making is not fully dependent on lightness (L*) of the sample *itself* as the animals were not systematically choosing darker or lighter samples in “same-colour” conditions.

When presented with two *green* samples, the winning choices tended to be more green and yellow. This may be related to the chlorophyll content, as shown in Table 3. However, no such differences existed when the animals had to choose between two red samples, where anthocyanin could be masking the chlorophyll pigments. The animals consistently chose the red samples containing greater amounts of anthocyanin despite there being no differences in the levels of chlorophyll and in a* or b* chromaticity channels (see fig. 6 & 7). Without further studies on differing plants containing differing amounts of anthocyanin we can only speculate on reasons for this. First, perhaps the red cabbage contained *too much* anthocyanin, and our failure to find any difference in RH/RL was because both groups exceeded a perceptual threshold in any reward system. Alternatively, perhaps visual differences do exist between the samples we chose, and our equipment did not detect this. However, this does not explain why they were able to pick the sample with greater amounts of anthocyanin. It is also important to note that colour measurements are not responsible for a unique set of pigments and can result from different combinations of pigments (22).

Overall, captive bred *Pongo pygmaeus* and *Pongo abelii* choose to eat samples of cabbage that contain a higher amount of anthocyanin and that their choice cannot be explained by a difference in size or lightness. Additionally, their choice can only be explained by a difference in colour when it comes to the green cabbage (*Brassica oleracea var.capitata*), but not the red (*Brassica oleracea var. capitata f. rubra*). Our second hypothesis then, that colour cues may guide orangutan choice in the absence of any odour, is only partially supported.

Traditionally anthocyanin, carotenoid and chlorophyll have been measured using destructive techniques but the development of non-destructive methods has allowed fast and simple analysis of plant pigments (63). The red and green cabbage leaves used in this study contained both anthocyanin and chlorophyll which were positively correlated, so it was important to be able to distinguish between these two pigments to resolve whether the Orangutans were selecting cabbage leaves based on levels of anthocyanin or chlorophyll. In a survey of 53 species, Sims and Gamon (2002) found no significant effect from either carotenoids or anthocyanins when measuring chlorophyll in vivo using spectral indices (64). Additionally the SPAD chlorophyll meter has previously been used to accurately detect levels of chlorophyll without interference from anthocyanin in young leaves of *Eucalyptus, Rosa, and Ricinus communis* (65). In contrast, Hlavinka, Nauš, & Špundová (2013) found that while the measured chlorophyll content was not significantly influenced by anthocyanin in mature green tomato leaves, in mature senescent leaves with SPAD readings below 20, anthocyanin substantially affected chlorophyll readings. As this study contained mature, non-senescent leaf material it is expected that increased levels of anthocyanin in the red cabbage was not contributing to an artificially elevated chlorophyll reading.

Moreover, the red and green cabbage leaves stored in the fridge for 24hrs prior to being presented to the orangutans showed a small increase in the levels of anthocyanin, yet they showed a small decrease in the levels of chlorophyll. A decrease in chlorophyll levels during storage is common in leafy vegetables (67). Similar results were reported in *valeriana* lettuce which showed a decrease in levels of chlorophyll over 5-7 day post-harvest but an increase in anthocyanin content over the same time period (68). Again, this points to a significant role for anthocyanin in the decision making task.

Anthocyanins are plant pigments frequently found in the human diet (69). They are the most commonly consumed plant secondary metabolites, with the richest sources coming from fruits and flowers (24). They are potent antioxidants that have been associated with roles in cardiovascular disease prevention and using in vitro cell based assays the anti-carcinogenic properties of anthocyanins has been well documented in; apoptosis (70), cell cycle arrest (71) and inhibition of DNA oxidative damage (72). Although conclusive clinical studies are limited, consumption of anthocyanins has been recommended as part of a “healthy heart” diet (73). Cognitive benefits have also been highlighted in studies on rats taking blueberry extract (37,38). The orangutans in the present study showed a strong preference for anthocyanin rich cabbage leaves in all 6 leaf combinations. This is in agreement with Schaefer et al., (2008) who found that wild European Blackcaps preferred anthocyanin rich fruits (42). They postulated that anthocyanin acts as a visual signal for total antioxidant content as it correlated with colourless flavonoid antioxidants such as flavones, isoflavones, flavonones, catechin, and isocatechin in fruit and berries. A strong correlation between anthocyanin and colourless flavonoids is also seen in cabbage (51). This is especially true in red cabbage which is unique among vegetables in containing a very high number of different anthocyanin compounds (49).

Our results build on these earlier studies, and as originally suggested by Schaefer et al., (2008) in their study on birds, we find that these trichromatic apes are capable of discriminating anthocyanin reward value in two types of cabbage without the ability to take advantage of the UV spectrum. Whether that reward signals for antioxidant content, anti-inflammatory properties or cognitive benefits is still to be established. Perhaps it is an example of self-medication by these animals (74)? Nevertheless, the reward is enough to influence *voluntary* decision making in these hominids. And, just as migrating juvenile blackcaps show colour preferences that differ for fruits and insects (32), perhaps the orangutans are also capable of differing colour preferences dependent upon the diet available at that time. They may be using different mechanisms when choosing between two reds than when choosing between two greens.

The captive animals in this study had daily access to a balanced diet of fruit and vegetables. Beyond the few enrichment activities provided by staff and volunteers they have no experience in foraging. They also have no knowledge of the various strategies that wild orangutans employ during times of low fruit abundance. Wild orangutans are omnivorous, feeding on fruit, seeds, bark, pith from palm and monocot stems, leaves, flowers, and insects such as termites and ants (75–78). They prefer fruit, but resort to bark, leaves and *fallback* foods during periods of low fruit availability (76). Examples of fallback foods containing large amounts of anthocyanin available to wild apes in Africa are fig fruits (*Ficus sansibarica*) (79,80), while in Asia species of *Neesia* are available to Orangutans (81,82). The nature and availability of fallback food is postulated to influence the socioecology and specific traits in apes such as harvesting adaptations or tool use (83,79) and perhaps, in addition to antioxidant properties and benefits mentioned above, rewards for consumption of anthocyanin by apes might be the strengthening of traits such as efficient foraging and harvesting (79) or advanced manual prehension needed to access the fruit at the *appropriate* stage of maturity (80).

In conclusion, learning any behavioural routine that facilitates survival is reinforced via rewards. Our animals quickly learnt the decision making task; voluntarily choosing samples with more anthocyanin. They did not make decisions based on lightness or red/purple. This is the first study to show this in non-human primates. Though we cannot extrapolate our findings to wild orang-utans without further study, we might speculate that anthocyanins reward for more than one beneficial action and that just as the *changing* value of a reward in fields such as economics or learning theory is important in many decision-making models, perhaps the value of anthocyanin as a reward changes as fruit becomes more abundant or ripe.

## Acknowledgments

We are grateful to all staff at the National Zoo of Malaysia (Zoo Negara) for their help on this project. We would particularly like to thank the zookeepers at the ape centre; Sharul, John and Firdaus for their assistance and endless patience. We would also like to acknowledge the skills of Danial Hafeez and Research Assistant Nadia Zulkilfi, in collecting and collating data. Many thanks to the Director, Zoology, Hospital & Veterinarian Services & Giant Panda Conservation Centre, Dr. Mat Naim Bin Haji Ramli. We also appreciate advice on enrichment from Zoo Curator, Doreen Khoo, and Education Officer, Junaidi Bin Omar. This research was possible through grants awarded from the Malaysian Ministry of Science, Technology and Innovation (MOSTI), grant number 06-02-12-SF0187, and from the Malaysian Ministry of Education, Fundamental Research Grant Scheme (FRGS) no. F0004.54.02.

References

